# Differential gene expression associated with a floral scent polymorphism in the evening primrose *Oenothera harringtonii* (Onagraceae)

**DOI:** 10.1101/2021.01.12.426409

**Authors:** Lindsey L. Bechen, Matthew G. Johnson, Geoffrey T. Broadhead, Rachel A. Levin, Rick P. Overson, Tania Jogesh, Jeremie B. Fant, Robert A. Raguso, Krissa A. Skogen, Norman J. Wickett

## Abstract

**Background:** Plant volatiles play an important role in both plant-pollinator and plant-herbivore interactions. Intraspecific polymorphisms in volatile production are ubiquitous, but studies that explore underlying differential gene expression are rare. *Oenothera harringtonii* populations are polymorphic in floral emission of the monoterpene (R)-(-)-linalool; some plants emit (R)-(-)-linalool (linalool+ plants) while others do not (linalool-plants). However, the genes associated with differential production of this floral volatile in *Oenothera* are unknown. We used RNA-Seq to broadly characterize differential gene expression involved in (R)-(-)-linalool biosynthesis. To identify genes that may be associated with the polymorphism for this trait, we used RNA-Seq to compare gene expression in six different *Oenothera harringtonii* tissues from each of three linalool+ and linalool-plants.

**Results:** Three clusters of differentially expressed genes were enriched for terpene synthase activity: two were characterized by tissue-specific upregulation and one by upregulation only in plants with flowers that produce (R)-(-)-linalool. A molecular phylogeny of all terpene synthases identified two putative (R)-(-)-linalool synthase transcripts in *Oenothera harringtonii*, a single allele of which is found exclusively in linalool+ plants.

**Conclusions:** By using a naturally occurring polymorphism and comparing different tissues, we were able to identify genes putatively involved in the biosynthesis of (R)-(-)-linalool. Expression of these genes in linalool-plants suggests a regulatory polymorphism, rather than a population-specific loss-of-function allele. Additional terpene biosynthesis-related genes that are up-regulated in plants that emit (R)-(-)-linalool may be associated with herbivore defense, suggesting a potential economy of scale between plant reproduction and defense.

## Background

Floral fragrance is a complex trait that is typically composed of tens to hundreds of volatile organic compounds (VOCs) whose ecological roles include the attraction of pollinators, manipulation of pollinator behavior, and defense against herbivores [1–7]. Considerable qualitative and quantitative variation in floral fragrance has been documented, with over 1700 floral volatiles having been described from more than 900 angiosperm species [8]. In addition to substantial interspecific variation, floral fragrance has also been shown to differ intra-specifically [2, 9]. The evolutionary and ecological mechanisms that maintain intraspecific variation in floral fragrance constitute an active area of research and have been attributed to differences in community composition, pollinator preferences, phenotypic plasticity, and genetic drift [10–13].

The biosynthetic pathways that produce floral volatiles are increasingly well-characterized [14], and recent studies have used transcriptomics-based approaches to identify homologous genes from these pathways in non-model plant species [15–18]. Despite these advances [19, 20], relatively few studies have identified the genetic mechanisms that impact how these pathways modify floral scent or how selection acts on the resulting intraspecific variation. Volatile terpenes comprise a major source of biosynthetic diversity in scents, primarily 10-carbon monoterpenes and 15-carbon sesquiterpenes [21]. Terpene synthase (TPS) enzymes, which are involved in the biosynthesis of these secondary metabolites, are encoded by a moderately large gene family in angiosperms [22]. The TPS genes exist in seven to eight subfamilies that show either lineage-specific (e.g. exclusively gymnosperm TPSs) or molecule/lineage-specific (e.g. monocot sesquiterpene TPSs) affinities. Despite the widespread characterization of the gene family and their products across plant lineages [22, 23], the mechanisms maintaining intraspecific variation in the production of volatile terpenes remain poorly understood. The functional implications of intraspecific terpenoid variation are epitomized by the culinary herb *Thymus vulgaris* (Lamiaceae), whose distinct “chemotypes” differ in herbivore defense, frost sensitivity, and allelopathic interactions with neighboring plants [24, 25]. Genetic crosses helped to document an epistatic network of mendelian loci responsible for volatile chemotypes in *T. vulgaris* [26, 27]. However, beyond a few such model systems, the genetic controls of chemotype variation remain largely unknown, and this knowledge is needed to understand how conflicting selective forces shape chemical polymorphism. To date, there have been few opportunities to study the genetic underpinnings of volatile terpenoid polymorphism in floral scent.

The focal species of this study, *Oenothera harringtonii* (Onagraceae), is a night-blooming, self-incompatible annual herb [28] that displays intraspecific variation among populations in the floral scent compound, (R)-(-)-linalool. Extensive pilot surveys (335 samples, 15-30 individuals per population, 12 populations) revealed that (R)-(-)-linalool is either emitted as an abundant floral volatile or is absent from the floral headspace of *O. harringtonii* plants (Additional File 4). Both enantiomers of this monoterpene alcohol, (R)-(-)-linalool and (S)-(+)-linalool, have been reported in a wide variety of plant families and are common components of floral scent, especially in nocturnal moth-pollinated flowers [29]. Other night-blooming species across the genus *Oenothera* have been found to emit (R)-(-)-linalool, including *O. biennis* (sect. *Oenothera*; [30]), *O. acutissima* (sect. *Lavauxia*; [31]), *O. californica* (sect. *Anogra*), *O. cespitosa* (sect. *Pachylophus*), *O. howardii* (sect. *Megapterium*), *O. lavandulifolia* (sect. *Calylophus*) and *O. xylocarpa* (sect. *Contortae*); Jogesh, Skogen, and Raguso unpublished data). Linalool in floral tissues can function as a pollinator attractant or as a defense compound [32–35], whereas linalool emitted by vegetative tissues has been implicated in both direct and indirect plant defenses [34]. Flowers of *O. harringtonii* are pollinated by the widespread hawkmoth species *Hyles lineata* and *Manduca quinquemaculata* [36]. Parallel studies suggest that these hawkmoths are likely attracted to the strong floral scent and high visual contrast of *O. harringtonii* flowers [37, 38]. In addition to its role in plant reproduction, the primary pollinator *H. lineata* is also an herbivore of the plant, with its caterpillars feeding on leaves and floral tissues. Furthermore, the developing seeds of *O. harringtonii* are consumed by caterpillars of the moth genus *Mompha* [39], whose adults oviposit on flowers at dusk [40]. Considering these selective forces, it is likely that both floral scent and volatiles emitted by vegetative tissues play important roles in the complex fitness landscape of this species. However, understanding how these selective forces mold volatile terpene variation requires the identification of genes, including (R)-(-)-linalool synthase and related terpene synthases, that are associated with scent polymorphism in *O. harringtonii*. In this way, identification and subsequent manipulation of the (S)-(+) linalool synthase gene in a wild tobacco, *Nicotiana attenuata*, led to novel insights about geographic variation in selective pressures and the importance of terpenoid volatile background in a tri-trophic/plant defense context [41].

Within the family Onagraceae, the gene encoding (S)-(+)-linalool synthase was identified from *Clarkia breweri* and *Clarkia concinna* in the first study of biosynthetic genes responsible for floral scent [42]. However, (R)-(-)-linalool synthase has not been identified in any evening primrose species to date, and only a single, partial TPS sequence has been identified thus far in *Oenothera* [43], one of the most species-rich lineages within the family [44]. In this study, we use RNA-Seq to identify differentially expressed genes (DEGs) correlated with linalool production in *O. harringtonii* by sequencing transcriptomes of six different tissues from three biological replicates of each of two chemotypes: (1) flowers that emit (R)-(-)-linalool (linalool+) and (2) flowers that do not emit (R)-(-)-linalool (linalool-). We use this information, in combination with a phylogenetic analysis of terpene synathases, to identify a gene that likely encodes (R)-(-)-linalool synthase in *O. harringtonii* and identify an allele that is specific to linalool+ individuals. This study includes the first transcriptomes assembled for *O. harringtonii*, providing critical genomic information to inform ongoing studies of intraspecific floral scent variation within this species. In addition, the preliminary identification of (R)-(-) linalool synthase and other differentially expressed transcripts is a first step towards a broader, comparative analysis of the molecular evolution of floral scent among species of *Oenothera* that differ in mating system, pollination mode, life history strategy and, thus, the direction and intensity of natural selection (see [45]). As such, results of the present study provide a key foundational step in understanding the ubiquity and relevance of (R)-(-)-linalool throughout flowering plants.

## Results and Discussion

### Floral Fragrance Analysis

Floral scent profiles were collected from replicated linalool+ and linalool-plants from which expressed transcripts were subsequently harvested; volatile chemical data are summarized in Table 1. Across all individuals, the floral scent was dominated by the monoterpene olefin (*E*)*-* β-ocimene, with small amounts of other monoterpenes ((*Z*)*-*β-ocimene and β-myrcene) and sesquiterpenes (β-caryophyllene and (*E,E*)-farnesol). The flowers of three individuals emitted large amounts of (R)-(-)-linalool (linalool+), whereas those from another three individuals did not emit any (Fig. 1A). This pattern is consistent with phenotypic data from the source populations for these plants (Additional File 4), confirming that linalool chemotypes breed true in greenhouse-grown plants (see Methods). Other volatiles (ocimene epoxide, methyl geranate, isophytol, α-humulene, and methyl benzoate) were emitted in small amounts by some individuals but not others, but the presence of these compounds varied independently of linalool chemotype (as they did in field-collected samples; Additional File 4) and are therefore likely due to unrelated factors.

**Table 1:** Volatile floral terpenoids and their emission rates for greenhouse-grown plants used for transcriptomics. Emission rates are given in μg per flower per hour. Volatile terpenoids are ordered by sub class (see Table S1) and retention time (min). The identity of all compounds except isophytol (*) and ocimene epoxide (*) was confirmed through co-chromatography and mass spectral comparison with synthetic standards.

|  | A | B | C | D | E | F |
| --- | --- | --- | --- | --- | --- | --- |
| **Monoterpenes** |  |  |  |  |  |  |
| β-myrcene | 0.18 | 0.15 | 0.13 | 0.07 | 0.03 | 0.09 |
| *cis*-β-ocimene | 0.62 | 0.54 | 0.39 | 0.19 | 0.09 | 0.27 |
| *trans*-β-ocimene | 27.50 | 22.49 | 23.26 | 14.08 | 9.28 | 19.67 |
| Ocimene epoxide* | 0.29 | 0.76 | - | - | 0.13 | 0.02 |
| **R-(-)-linalool** | **15.25** | **13.79** | **12.23** | - | - | - |
| **Diterpenes** |  |  |  |  |  |  |
| Isophytol* | 0.04 | 0.08 | - | - | - | - |
| **Sesquiterpenes** |  |  |  |  |  |  |
| β-caryophyllene | 0.42 | 0.09 | 0.05 | 0.03 | 0.46 | 0.02 |
| α-humulene | 0.03 | - | - | - | 0.01 | - |
| *trans-trans*-Farnesol | 1.07 | 0.23 | 0.03 | 0.02 | 0.05 | 0.02 |
| Methyl farnesoate | 6.79 | 0.70 | 0.07 | - | 0.54 | 0.02 |

**Figure 1:**
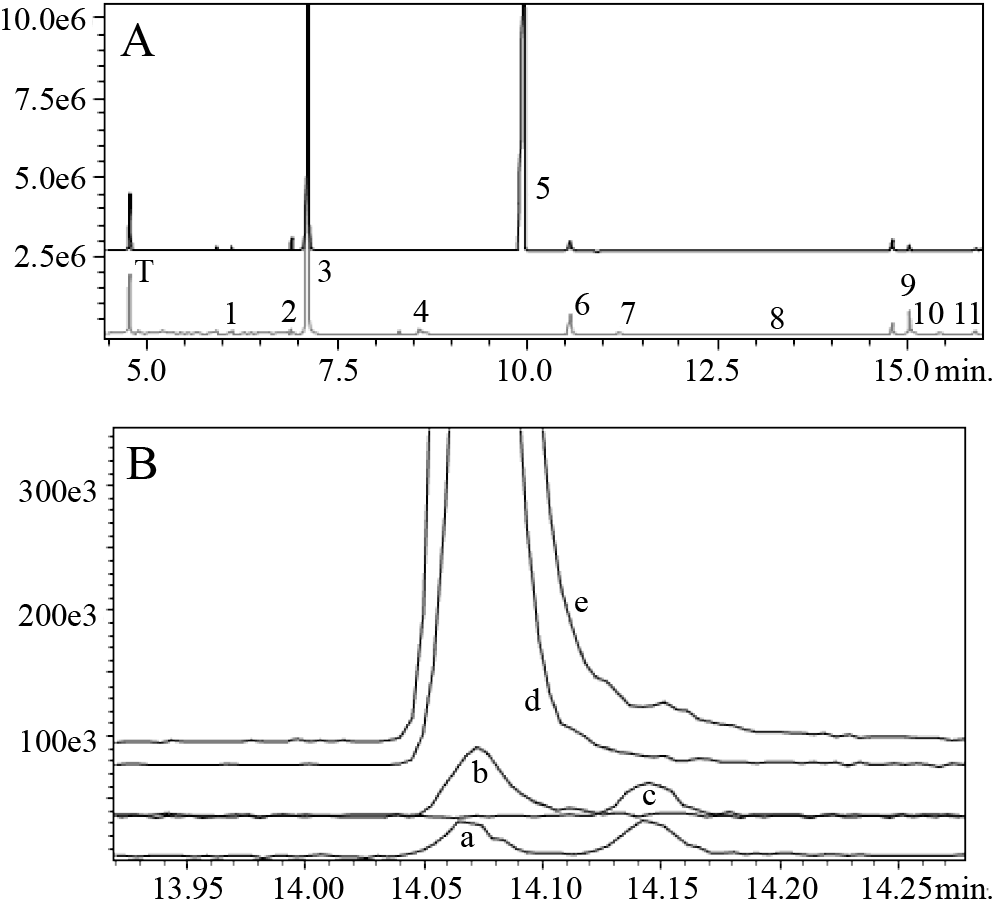
Gas chromatographic representation of differences between chemotypes of *Oenothera harringtonii*. A. Total ion chromatogram of floral scent from two accessions, the upper trace (black) from a linalool+ plant and the lower trace (gray) from a plant lacking this compound. Other than the internal standard (toluene; T), peak numbers indicate floral scent compounds in the order listed in Table 1 (e.g. 5: linalool), with retention times given for a typical polar (EC-wax) GC column. B. Separation of linalool enantiomers on a chiral GC column, confirming that flowers of *O. harringtonii* exclusively emit (R)(-) linalool. Trace a = racemic (1:1) (R)(-) and (S)(+) linalool, trace b = synthetic (R)(-) linalool, trace c = (S)(+) linalool in floral headspace of *Clarkia breweri*, trace d = floral headspace of *O. harringtonii* from Florence, Colorado (FLO), and trace e is the same sample spiked with (R)(-)linalool. In both panels, vertical axis shows counts (abundance) and horizontal axis shows retention time (minutes).

Additional analyses were performed to confirm which of the two enantiomers of linalool were present in the floral scent. Chiral GC-MS analyses resulted in the separation of linalool enantiomers to baseline, with (R)-(-)-linalool (14.07 min) eluting before (S)-(+) linalool (14.14 min) under the conditions described (Fig. 1B). When the linalool peaks from linalool+ populations (Florence, Maverick, Baculite Mesa) were found to align with the retention time of the (R)-(-) enantiomer, but not with that of the (S)-(+) enantiomer or the linalool peak from *Clarkia breweri*, we re-injected an *O. harringtonii* headspace sample from the Florence population, to which a small amount of this enantiomer was added. These analyses confirmed that flowers of *O. harringtonii* exclusively emit (R)-(-) linalool (Fig. 1B).

### Transcriptome Sequencing, Assembly, and Annotation

RNA was successfully sequenced from six tissues (stamens, stigma/style, petals, hypanthium, bud, and leaf) for all individuals, except for individuals D and F, for which bud and leaf samples were not obtained (see Methods). Across all 32 samples, 1.57 billion paired reads were sequenced (Additional file 1). On average, 95.77% of reads passed quality trimming, and 99.04% of these high-quality reads were retained after filtering for plastid genes and rRNA. In total, 94.86%, or 1.49 billion reads, were retained after filtering. After normalization, a total of 33 million reads were assembled into a reference assembly consisting of 489,895 contigs and 244,284 components. Through alignment and abundance estimation using RSEM, 72% of cleaned reads aligned to the reference assembly. Median contig length was 393 base pairs, average contig length was 676.8 base pairs, and transcript N50 was 1047 base pairs. As determined using the Trinotate pipeline [46], 27.6% of contigs had either a blastx or blastp hit to Swiss-Prot, and 25.5% of contigs had a gene ontology term assigned from either blast or pfam. In a blastx search of the assembly against an *Arabidopsis* database [47], 174,068 contigs had a hit in the *Arabidopsis thaliana* protein database; 20,480 of these were unique *Arabidopsis* protein hits. In a tBLASTn search of *Arabidopsis* proteins against the assembly, 31064 *Arabidopsis* peptides had a hit and 14,614 of these were unique. In total, there were 11,754 reciprocal best hits between the transcriptome assembly and the *Arabidopsis* proteome, with an ortholog hit ratio of 0.34 and an average collapse factor of 2.13.

### Differential Expression and GO Enrichment Analysis

Principal components analysis was conducted to test whether it is appropriate to group biological replicates together during subsequent differential expression analysis. In general, biological replicates of the same tissue and chemotype clustered together (Fig. 2). Additionally, samples of the same linalool chemotype grouped together, suggesting significant gene expression differences between the chemotypes. In total, edgeR identified 6,577 differentially expressed genes (DEGs) among the six tissues and between linalool+ and linalool-individuals, with an overall pattern suggesting that tissue identity explains most of the observed differential expression (Fig. 3). All DEGs were grouped into 12 clusters by cutting the DEG dendrogram at 65% of its height (Fig. 4). Gene ontology (GO) enrichment analysis was performed for each cluster to identify categories of genes that are overrepresented (see Additional file 2 for a complete list of enriched GO terms for each cluster).

**Figure 2:**
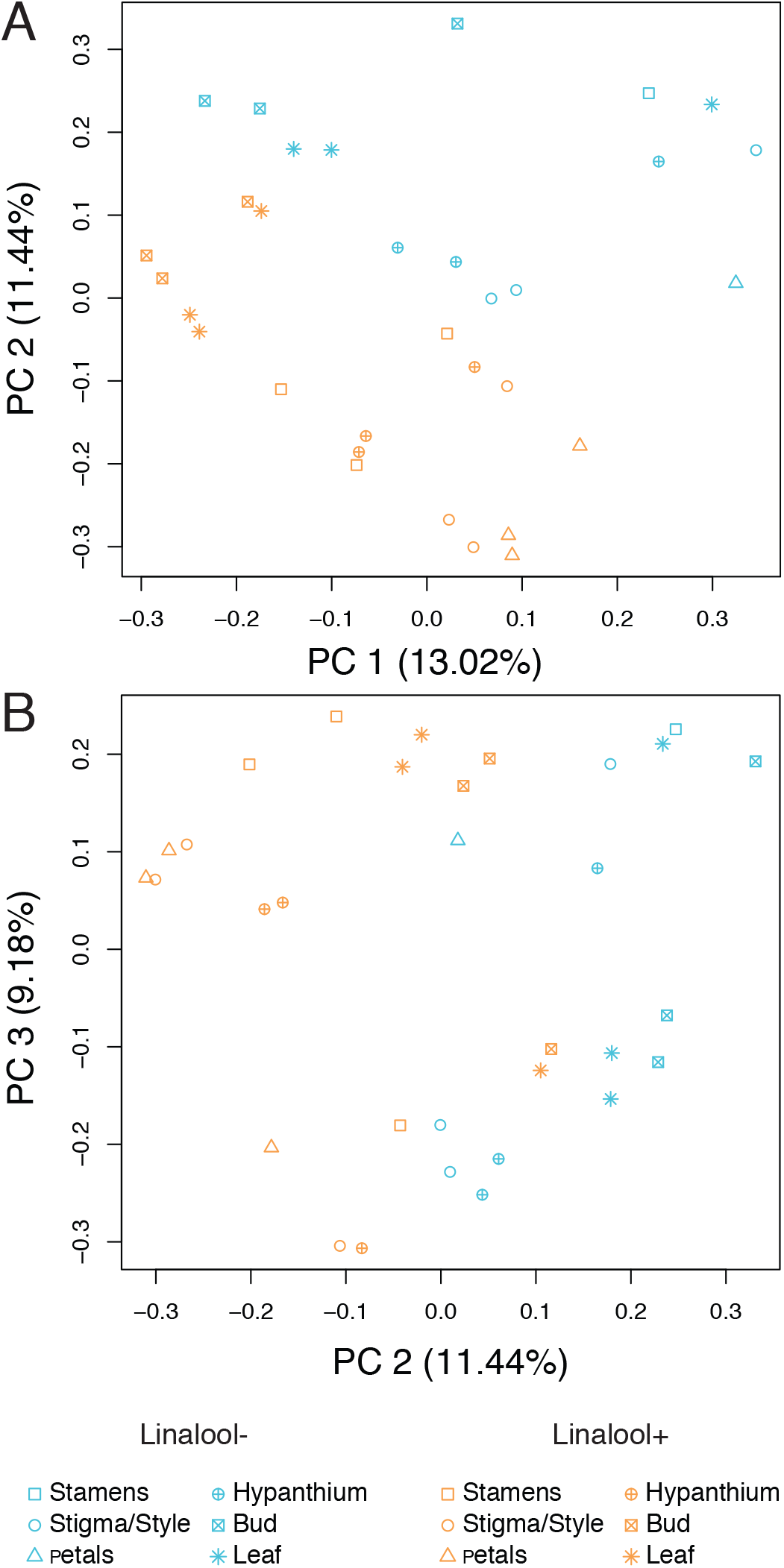
Principal Components Analysis of Differential Gene Expression. Each RNA-Seq library is represented by a symbol whose shape indicates tissue source and color indicates linalool chemotype (blue = linalool- and orange = linalool+). Overall, 33.64% of variation in gene expression is captured by the first three principal components. (A) PC 1 and 2 (B) PC 2 and 3.

**Figure 3:**
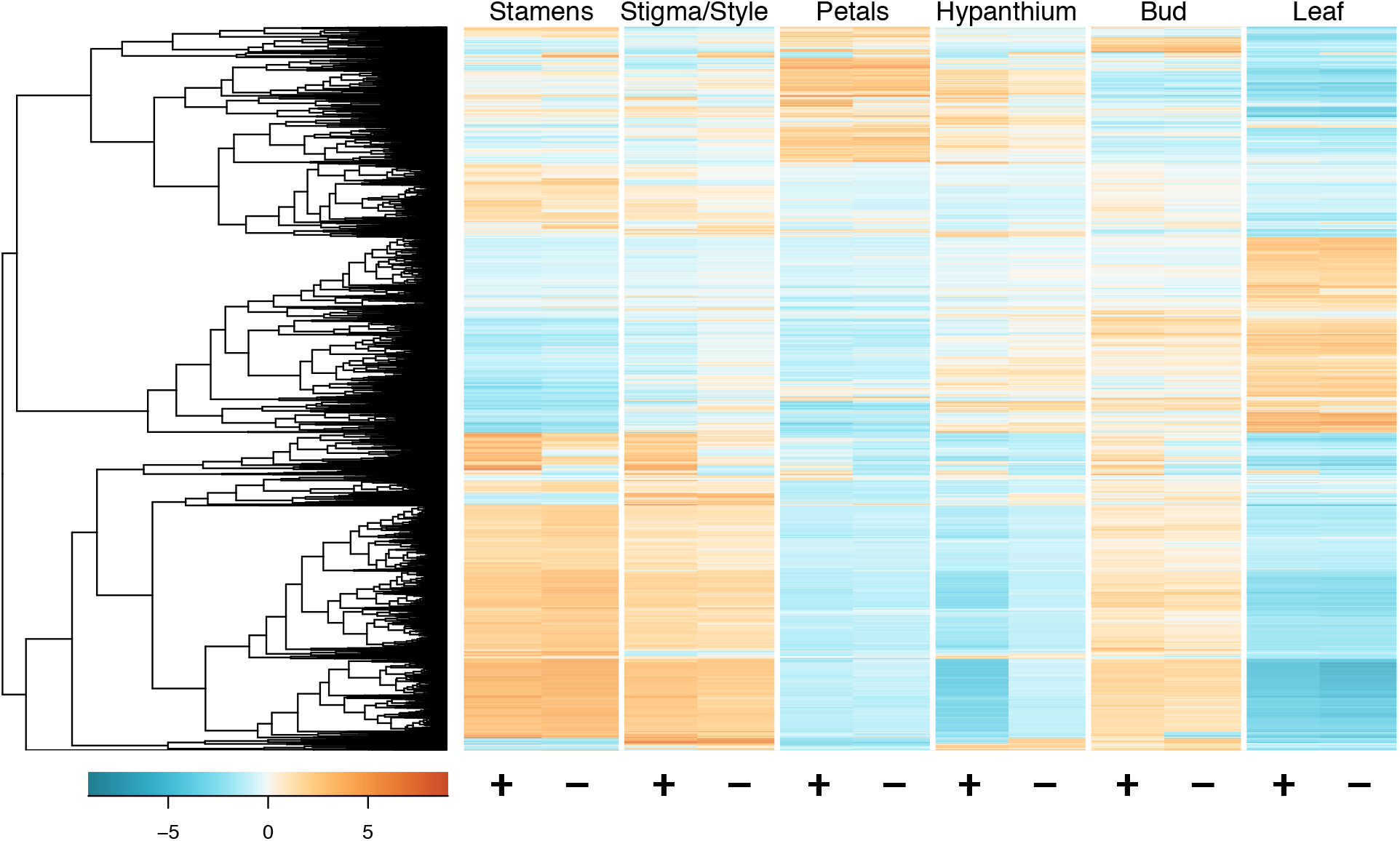
Heatmap of Expression for all Differentially Expressed Genes. In total, 6577 genes were identified as differentially expressed. Biological replicates are pooled together; orange coloration indicates upregulation of genes and blue indicates downregulation. Genes are ordered by similarity of expression, as represented by the dendrogram on the left side of the heatmap.

**Figure 4:**
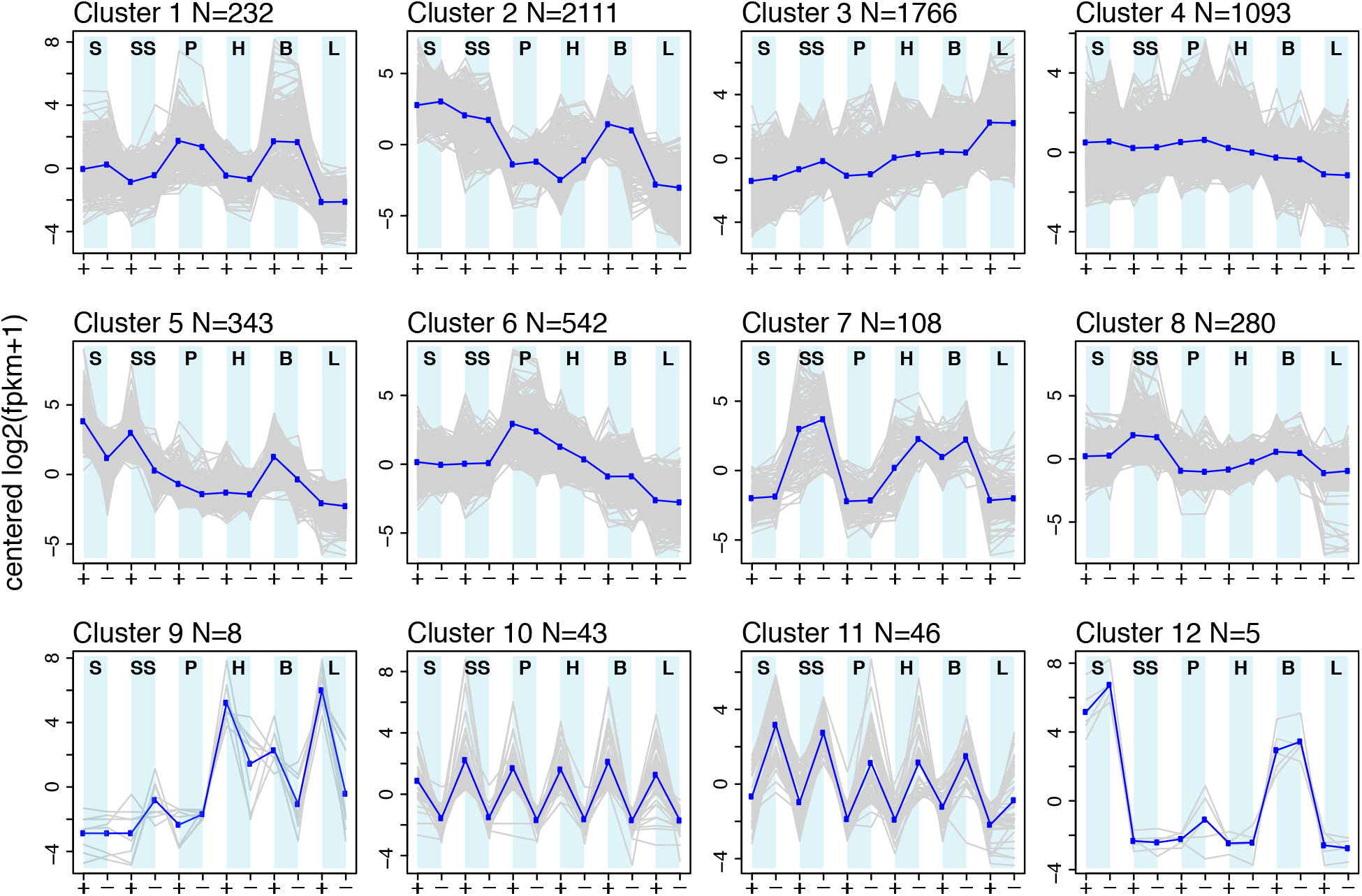
Expression Patterns for Clusters of Differentially Expressed Genes. Each line graph represents relative expression for a group of differentially expressed genes with similar expression patterns, as defined by cutting the dendrogram in Figure 3 at 65% of its height. Grey lines indicate expression of individual genes and dark blue lines indicate average overall expression pattern for that cluster. S = stamens, SS = stigma and style, P = petals, H = hypanthium, B = bud, and L = leaf.

Of the twelve clusters, two (clusters 10 and 11) were characterized by differential expression of genes between linalool+ and linalool-individuals, suggesting that genes in these clusters may be involved in linalool biosynthesis. Genes in cluster 10 (n=43) were upregulated in all tissues from linalool+ individuals, whereas genes within cluster 11 (n=46) were downregulated in all tissues from linalool+ individuals. In general, both of these clusters were characterized by several genes that lack annotations in the Trinotate report (26/43 and 38/46 genes for each cluster, respectively), either because they are unique or significantly differ from a homolog in the reference database.

To obtain an overview of the functions of genes within each cluster, we conducted GO enrichment. Cluster 10 is enriched for terpene synthase activity (Table 2), and transcripts in this cluster with this higher-level GO term (TPS activity) include those annotated as terpene synthase 10 (DN74292_c2_g1), γ-terpinene synthase/(+)-α-terpineol synthase (TRINITY_DN62538_c3_g2) and santalene synthase (DN76019_c5_g5). Terpene synthase 10 is known to produce (R)-(-)-linalool as a primary product in *Arabidopsis thaliana*, with β-myrcene and (*E*)-β-ocimene as minor products [48]. γ-terpinene and (+)-α-terpineol are monoterpenes, and although santalene is a sesquiterpene, terpene synthases responsible for its biosynthesis are classified with the TPS-b monoterpene synthases [49]. γ-terpinene and santalene are not present in the floral scent of *O. harringtonii*, whereas α-terpineol is only present in small amounts, inconsistently, in some populations (e.g. FLO; Additional File 4). Thus, either the appropriate biosynthetic enzymes are not translated from these transcripts or, if they are, their eventual products are either metabolized into other secondary compounds, or they simply are not volatilized. These results suggest that linalool+ individuals have the capacity to synthesize terpenes in all six tissues included here; therefore, differences between the linalool+ and linalool-chemotypes include terpene synthase expression in both floral and vegetative tissues. In addition to being common components of floral fragrance, terpenes are known to be involved in constitutive and induced defenses in both floral and non-floral tissues [4, 50]. Terpenes are also involved in indirect defense, as they can attract members of the third trophic level [51–55]. Overexpression of TPS genes across all tissues including leaves suggests that *O. harringtonii* populations emitting R-(-) linalool from flowers may also be better-defended.

**Table 2:** GO Enrichments for each cluster. Top five significant enrichments (FDR>0.05) for each cluster are shown. Note that clusters 5, 10, 11, and 12 had fewer than five significant enrichments. Full list of enriched GO terms can be found in Additional File 2.

| Cluster | GO Term ID | Ontology | GO Term | FDR |
| --- | --- | --- | --- | --- |
| 1 | GO:0008610 | BP | lipid biosynthetic process | 1.68E-09 |
| 1 | GO:0044255 | BP | cellular lipid metabolic process | 1.14E-08 |
| 1 | GO:0008299 | BP | isoprenoid biosynthetic process | 1.85E-07 |
| 1 | GO:0019288 | BP | isopentenyl diphosphate biosynthetic process, methylerythritol 4-phosphate pathway | 1.91E-07 |
| 1 | GO:0046577 | MF | long-chain-alcohol oxidase activity | 2.69E-07 |
| 2 | GO:0030599 | MF | pectinesterase activity | 9.04E-66 |
| 2 | GO:0045490 | BP | pectin catabolic process | 1.56E-60 |
| 2 | GO:0045330 | MF | aspartyl esterase activity | 1.92E-56 |
| 2 | GO:0010393 | BP | galacturonan metabolic process | 1.30E-50 |
| 2 | GO:0045488 | BP | pectin metabolic process | 1.30E-50 |
| 3 | GO:0044435 | CC | plastid part | 7.58E-70 |
| 3 | GO:0044434 | CC | chloroplast part | 7.58E-70 |
| 3 | GO:0009579 | CC | thylakoid | 2.78E-62 |
| 3 | GO:0044436 | CC | thylakoid part | 5.12E-53 |
| 3 | GO:0009507 | CC | chloroplast | 1.85E-51 |
| 4 | GO:0016054 | BP | organic acid catabolic process | 3.47E-09 |
| 4 | GO:0046395 | BP | carboxylic acid catabolic process | 3.47E-09 |
| 4 | GO:0009702 | MF | L-arabinokinase activity | 1.46E-08 |
| 4 | GO:0044282 | BP | small molecule catabolic process | 1.64E-07 |
| 4 | GO:1901606 | BP | alpha-amino acid catabolic process | 9.71E-07 |
| 5 |  |  | no significant terms |  |
| 6 | GO:0010333 | MF | terpene synthase activity | 4.03E-10 |
| 6 | GO:0016838 | MF | carbon-oxygen lyase activity, acting on phosphates | 1.00E-09 |
| 6 | GO:0010279 | MF | indole-3-acetic acid amido synthetase activity | 5.93E-08 |
| 6 | GO:0010334 | MF | sesquiterpene synthase activity | 2.57E-06 |
| 6 | GO:0008152 | BP | metabolic process | 1.17E-05 |
| 7 | GO:0005576 | CC | extracellular region | 3.96E-05 |
| 7 | GO:0000272 | BP | polysaccharide catabolic process | 8.36E-05 |
| 7 | GO:0045490 | BP | pectin catabolic process | 1.13E-04 |
| 7 | GO:0010393 | BP | galacturonan metabolic process | 1.45E-03 |
| 7 | GO:0045488 | BP | pectin metabolic process | 1.45E-03 |
| 8 | GO:0080054 | MF | low-affinity nitrate transmembrane transporter activity | 2.18E-04 |
| 8 | GO:0080055 | BP | low-affinity nitrate transport | 2.18E-04 |
| 8 | GO:0015706 | BP | nitrate transport | 7.68E-02 |
| 8 | GO:0015112 | MF | nitrate transmembrane transporter activity | 7.68E-02 |
| 8 | GO:0015833 | BP | peptide transport | 7.68E-02 |
| 9 | GO:0000272 | BP | polysaccharide catabolic process | 0 |
| 9 | GO:0004553 | MF | hydrolase activity, hydrolyzing O-glycosyl compounds | 0 |
| 9 | GO:0004568 | MF | chitinase activity | 0 |
| 9 | GO:0005976 | BP | polysaccharide metabolic process | 0 |
| 9 | GO:0006022 | BP | aminoglycan metabolic process | 0 |
| 10 | GO:0010333 | MF | terpene synthase activity | 1.83E-02 |
| 10 | GO:0016838 | MF | carbon-oxygen lyase activity, acting on phosphates | 1.83E-02 |
| 11 |  |  | no significant terms |  |
| 12 | GO:0006869 | BP | lipid transport | 2.77E-02 |

Cluster 11 did not have any significant GO term enrichments, possibly because only a small number of the transcripts in the cluster were able to be annotated. The few genes in cluster 11 that do have annotations are not associated with terpene biosynthesis (Additional File 2). Two additional clusters contained significantly enriched GO terms related to terpene biosynthesis: clusters 1 (n=232) and 6 (n=542). Cluster 1 comprised genes upregulated in buds and petals but downregulated in leaves. Significant (FDR > 0.05) GO term enrichments for this cluster include isoprenoid biosynthetic process, terpenoid biosynthetic process, and isopentenyl diphosphate biosynthetic process, among others (Table 2 and Additional file 2). Cluster 6 comprised genes upregulated in petals and downregulated in leaves. Significant GO term enrichments for this cluster include terpene synthase activity, sesquiterpene synthase activity, isoprenoid biosynthetic process, and terpenoid biosynthetic process (Table 2 and Additional file 2). Neither cluster showed substantial differentiation between the two linalool chemotypes, which, in combination with the enriched GO terms for each cluster, is an indication that the genes contained within those clusters may be involved in the biosynthesis of floral scent volatiles that are shared between the two chemotypes. Thus, the expression patterns of these two clusters show that certain terpenoid scent compounds are likely to be produced primarily in the petals of *O. harringtonii*, and that buds may be synthesizing floral scent or floral scent precursors 24-48 hours prior to anthesis.

### Phylogenomic Analysis of Terpene Synthases and (R)-(-)-linalool Polymorphisms

To further characterize and identify terpene synthases in the transcriptome, regardless of their expression pattern, a profile Hidden Markov Model (pHMM) for terpene synthase (TPS) proteins was created based on a multiple sequence alignment of TPS sequences from van Schie et al. [23]. The pHMM successfully identified 40 TPS inferred protein sequences in the *O. harringtonii* reference assembly (Fig. 5). While it is likely that this is an overestimate due to the co-assembly of multiple individuals for the reference transcriptome, the 40 sequences were successfully placed into five of the seven TPS subfamilies with high confidence. According to the TPS subfamily circumscription in van Schie et al. [23], we found: (1) two distinct sequences of diterpene synthases (TPSc); (2) a single sequence of (S)-(+)-linalool synthase (TPSf); (3) 11 sequences of a second group of diterpene synthases (TPSe) with seven distinct groups of genes (components arising from the Trinity assembly); (4) eight sequences of sesquiterpene synthases (TPSa) with six distinct components, and (5) 18 sequences of monoterpene synthases (TPSb). No sequences were reconstructed within the TPSd or TPSg subfamilies (Fig. 5); the former (TPSd) comprises exclusively gymnosperm terpene synthases and the latter (TPSg) comprises a group of acyclic monoterpene synthases related to ocimene synthases in *Antirrhinum* [23].

**Figure 5:**
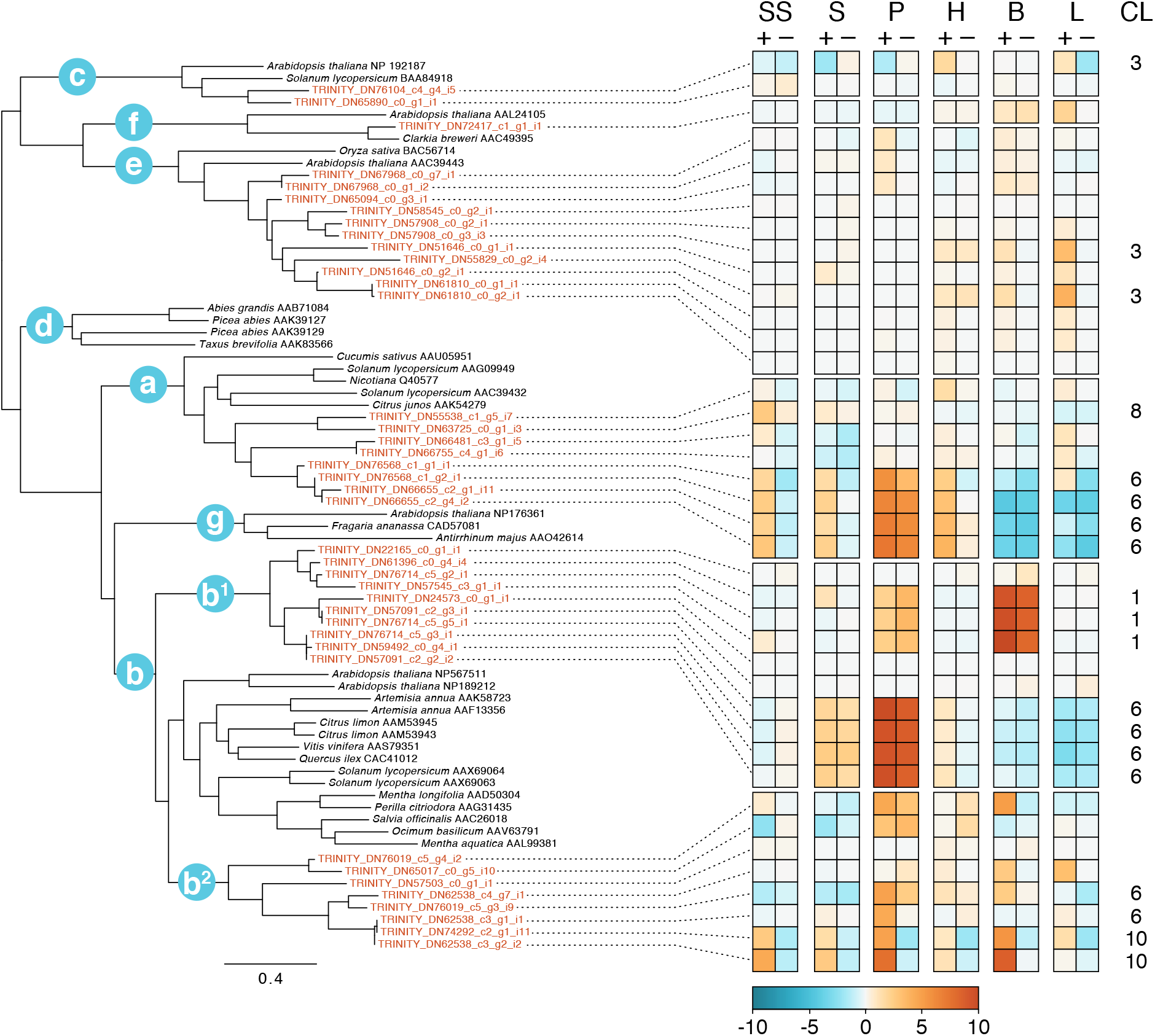
Terpene Synthase Gene Family Phylogeny and Gene Expression. Gene labels in red are putative *O. harringtonii* terpene synthases as identified by a pHMM based on sequences from van Shie et al. 2007, which are shown in black. The seven TPS subfamilies a-g are indicated on the tree; note that *O. harringtonii* transcripts fall into two clades within subfamily b, represented by b^1^ and b^2^. All TPS subfamilies are supported by BS > 95; full support values can be found in supporting materials. Relative expression for tissue-specific libraries (SS = stigma/style; S = stamens; P = petals; H = hypanthium; B = bud; L = leaf) for each *O. harringtonii* transcript is shown on the right, including its cluster number (CL) if the gene was differentially expressed.

Genes belonging to TPSc, TPSe, and TPSf largely showed no differential expression between either tissue or chemotype (Fig. 5). Two genes within the TPSe clade belong to DEG cluster 3 (Fig. 4 and 5) and were upregulated in leaves, suggesting that they may play a role in defense against herbivory rather than floral fragrance. Some sesquiterpene synthase genes (the TPSa clade) were upregulated in floral tissues of linalool+ individuals (see cluster 6 in Fig. 4; Fig. 5); however, these genes were highly expressed in both linalool+ and linalool-individuals in the petals. This matches fragrance data collected from source populations; both linalool+ and linalool-populations of *O. harringtonii* emit a range of sesquiterpenes that include β-caryophyllene, α-humulene, α- and β-farnesenes (Additional File 4).

To-date, the most intensive genetic and physiological study of floral fragrance emission in Onagraceae has focused on a pair of sibling species in the genus *Clarkia* [42, 43, 56, 57]. Most *Clarkia* species are bee-pollinated or autogamous [58], and range from unscented to mildly scented, with the exception of *C. breweri*, which produces a strong floral fragrance rich in (S)- (+)-linalool as a derived trait associated with a shift to hawkmoth pollination [57]. Pichersky et al. [56] found that (S)-(+)-linalool is primarily emitted from the petals and, to a lesser extent, the pistil of *C. breweri* flowers, whereas comparative analyses of the closest relative, *C. concinna*, revealed linalool production only in stigmatic tissues. The authors inferred that increased levels of linalool synthase (LIS) gene expression, expansion of tissues in which it is expressed, and enzymatic upregulation all contributed to the evolutionary gain-of-function of strong (S)-(+)-linalool emission from the flowers of *C. breweri* [29]. Our phylogenomic analysis placed a single sequence of (S)-(+)-linalool synthase (TPSf) from *O. harringtonii* as sister to (S)-(+)-linalool synthase from *Clarkia* (Fig. 5). While we cannot confirm that this putative (S)-(+) - linalool synthase in *O. harringtonii* is non-functional, it is clear that no detectable (S)-(+)-linalool is emitted from the flowers (Fig. 1B; Table 1), the transcript is not differentially expressed, and it is expressed at a low level.

The *O. harringtonii* genes circumscribed as belonging to the TPSb (monoterpene synthases) clade can be divided into two subclades, annotated here as TPSb^1^ and TPSb^2^ (Fig. 5). While both clades are strongly supported, their relationships to other TPSb genes from other species are not well resolved. Genes from *O. harringtonii* within the TPSb^1^ clade show some evidence of upregulation in floral tissues and bud (clusters 1 and 6) with no evidence of chemotype-related differential expression. As the two chemotypes differ primarily in the presence of a single monoterpene, (R)-(-)-linalool, we would expect that only one of the TPSb subclades would include a gene that is upregulated in linalool+ floral tissues. Two contigs from clade TPSb^2^ belong to cluster 10, which showed strong differential expression related to chemotype (see rows annotated as cluster 10 in Figure 5). Although cluster 10 includes genes that are differentially expressed (upregulated in linalool+ individuals) for all tissues including leaves, the signal for upregulation of these two transcripts in leaves is relatively weak. In fact, we detect no (R)-(-)-linalool emitted from the leaves of individuals with linalool+ flowers (Additional file 3). Furthermore, the TPSb^2^ subclade is sister (with moderate bootstrap support of 68) to a clade that includes other linalool synthases, including the (R)-(-)-linalool synthase gene from *Artemesia annua* [59]. When the phylogeny, differential expression, and floral fragrance data are taken together, they strongly suggest that the two transcripts belonging to cluster 10 in clade TPSb^2^ are (R)-(-)-linalool synthases in *O. harringtonii*.

Accounting for possible transcriptome assembly artifacts, we examined the three transcripts from the clade containing the candidate (R)-(-)-linalool synthases (DN62538_c3_g1_i1, DN62538_c3_g2_i2, and DN74292_c2_g1_i11) and determined that they each represent a partial coding sequence; the three transcripts were subsequently merged to form a putative full-length transcript. A BLAST search of the merged transcript revealed sequence similarity to (R)-(-)-linalool synthase in lemon myrtle, *Backhousia citriodora* (Myrtaceae; GenBank accession AB438045.1). However, the *O. harringtonii* transcript could only be aligned over 52% of the lemon myrtle transcript indicating that the merged *O. harringtonii* transcript may either not represent a full-length transcript, is shorter than the coding sequence in *B. citriodora*, or is significantly diverged in sequence (sequence similarity to the *B. citriodora* CDS is 64%). Further sequencing and functional analysis would be needed to determine which is the case. As RNA-Seq has been shown to reliably predict genes that are significantly associated with a treatment [60–63] and given the overwhelming difference between the abundance of reads mapped to the candidate (R)-(-)-linalool synthase transcripts in linalool+ and linalool-individuals, we suggest that RT-qPCR validation is unnecessary in this case. Future studies that focus exclusively on the function of this particular gene in *Oenothera* in more detail may benefit from the sequences described here, as a template for primer design or as a candidate locus in association studies.

To investigate if there are differences in the coding region of (R)-(-)-linalool synthase associated with either chemotype, we compared SNPs between individuals. Calling SNPs for each individual revealed that all linalool+ individuals are homozygous and share the same haplotype which matched the reference (merged consensus) sequence, with the exception of two SNPs at the 5’ and 3’ ends of the coding region (Fig. 6), which may be due to sequencing or assembly error (regions where read depth decreases). Conversely, all linalool-individuals are heterozygous, and do not share the same haplotype as each other or the linalool+ individuals. Sequences reconstructed from the linalool-individuals are characterized by between eight and fifteen SNPs compared to the reference sequence. The presence of a single allele in all linalool+ individuals included here is suggestive of a selective sweep in populations dominated by linalool+ plants, but much more extensive sampling of individuals and genomic regions is needed to better characterize the history of selection. Phasing of the alleles for linalool-individuals, and a reconstruction of the terpene synthase gene tree with additional sequences added (Additional File 5) confirmed that the sequences were indeed alleles rather than paralogs.

**Figure 6:**
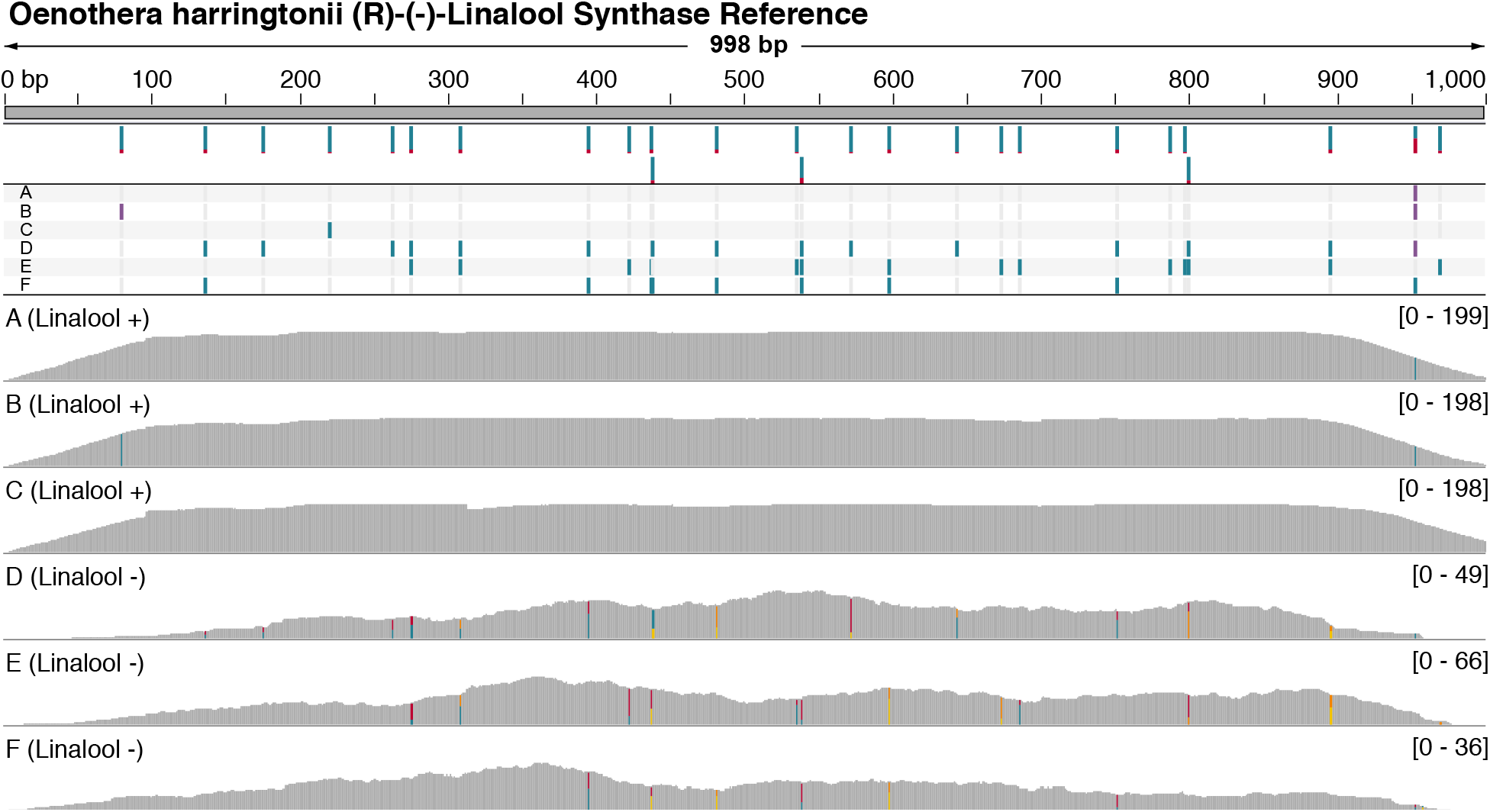
Distribution of polymorphic sites relative to a reconstructed (R)-(-)-Linalool Synthase reference gene. Top: Reference sequence and locations of SNPs passing quality filters. Colors indicate allele frequencies across the six individuals of Oenothera harringtonii. Middle: SNP calls passing filters for six individuals of Oenothera harringtonii relative to the reference sequence. Bottom: Depth of sequencing across the reference sequence for each individual (range of depth indicated to the right). Colors indicate presence of heterozygosity. Only deduplicated transcriptome reads were used for this analysis.

Terpenes are known to be regulated both transcriptionally and post-transcriptionally. In *Arabidopsis*, for example, differences in the amount of (*E*)-β-ocimene and (*E,E*)-α-farnesene emitted between two ecotypes is controlled by allelic variation of two terpene synthase genes, whereby one of the two genes in each ecotype produces a non-functional transcript [64]. Transcription factors regulating a variety of terpenes have also been identified; in cotton WRKY1 positively regulates (+)-□-cadinene synthase (CAD1) [65], and in kiwifruit, NAC transcription factors were found to regulate the expression of TPS1 in ripe fruit [66]. Post-transcriptional regulation includes the action of miRNAs, which have been identified in plants such as *Ocimum basilicum* [67] and *Ferula gummosa* [68]. The existence of SNPs between transcripts from each chemotype may suggest differences in functionality of each linalool synthase; however, few of the SNPs are shared among linalool-plants, such that the regulation of (R)-(-)-linalool synthase in *O. harringtonii* may be associated only with upregulation and the allele found in linalool+ individuals. Further sequence analysis and more extensive sampling is required to determine whether this apparent allelic diversity shows strong evidence of population structure. Additionally, a high-quality draft genome sequence for *O. harringtonii* would allow us to better detect variation that may be associated with regulatory regions or copy number variation.

## Conclusions

This study presents the first transcriptome assembly and analysis of differential expression in *Oenothera harringtonii*, a species with intraspecific variation for the emission of (R)-(-)-linalool in its floral fragrance. In this study, RNA-Seq was used to compare transcriptomes of six different *Oenothera harringtonii* tissues from six individuals (three linalool+, three linalool-). 6577 differentially expressed genes were identified, and these DEGs were sorted into 12 clusters. Three clusters were significantly enriched for GO terms related to floral fragrance biosynthesis. Two of these clusters showed upregulation in petals, suggesting that floral scent biosynthesis is mostly localized to petals. The third cluster showed upregulation in all linalool+ tissues and contained a significantly enriched GO term for terpene synthase. A phylogenetic analysis of all terpene synthases identified using a profile Hidden Markov Model classified 40 *O. harringtonii* transcripts into five terpene synthase subfamilies. Synthesizing the phylogenetic, differential expression, and floral fragrance data, we identified a transcript that likely encodes a partial (R)-(-)-linalool synthase, representing the first characterization of this gene in the flowering plant family Onagraceae.

## Methods

### Plant Material

Parental individuals were grown from seed collected in natural populations of *Oenothera harringtonii* from southeastern Colorado for which floral volatile data (Additional File 4) indicated that floral scent was polymorphic among the sampled populations: Burnt Mill Road (Pueblo County; linalool+), Florence (Fremont County; linalool+), Bloom (Las Animas County; linalool-), and David Canyon (Otero County; linalool-) (see [69] for locality data). As part of ongoing work on this species, the following controlled crosses within chemotype were conducted in the greenhouse at the Chicago Botanic Garden (Glencoe, Illinois, USA) in 2008 and 2009: both parents from Burnt Mill Road (individuals A and B), both parents from Florence (individual C), one parent from David Canyon and the other from Bloom (individual D), and both parents from David Canyon (individuals E and F). The resultant offspring seeds were germinated and grown in the greenhouse until buds were initiated, after which they were moved to a growth chamber (Conviron model GR48, Winnipeg, MB, Canada) where they typically flowered within one week. While anthesis in natural populations occurs at dusk [69], to facilitate daytime floral fragrance sampling, the growth chamber settings were configured such that dusk began at 9:30, night was from 11:00 to 20:30, dawn began at 20:30, and daytime was from 21:30 to 9:30.

### Floral Fragrance Collection

Based on hundreds of previous analyses of floral scent from across the range of *O. harringtonii* (Additional File 4), the population-level chemotypes are stable and consistent; floral scent analyses of the plants grown for this transcriptomic study were carried out to confirm the prediction of chemotypes for the plant material selected for RNA-Seq. Floral fragrance was collected prior to RNA extraction to confirm the linalool chemotype (linalool+ or linalool-) of each of the six individuals. Each flower was enclosed in a nylon resin oven bag (Reynolds, Lake Forest, IL) that was cut to 20.3cm x 14.8cm and resealed. Filters, constructed from a glass Pasteur pipette containing 10 mg of adsorbent material (Porapak Q 80/100 mesh size, Sigma-Aldrich, St. Louis, MO) tamped between quartz wool, were used to trap floral volatiles during the first hour after anthesis at dusk. The bag was secured around the flower’s hypanthium and the filter with twist-ties, ensuring that there were no vegetative structures enclosed within the bag.

The filter tip was positioned as close as possible to the floral center without damaging it and attached via Teflon tubing to a vacuum pump (PAS-500, Spectrex, Redwood City, CA). Pumps were run at a flow rate of 200ml air per minute for one hour; filters were then eluted with 200μl of high purity hexane (GC2, Burdick & Jackson, Muskegon, MI). Two types of controls were used for each floral fragrance collection: an ambient control filter attached to a bag without a flower, and a clean storage control filter that was eluted without having been used for fragrance collection. Eluted samples were stored at −20°C prior to being analyzed using gas chromatography-mass spectrometry (GC-MS) at Cornell University.

### Floral Scent Analysis by GC-MS

Prior to analysis, hexane-eluted floral headspace samples were concentrated to a uniform volume of 50μl using gaseous N_2_, and 5μl of 0.03% toluene in hexane (23ng) was added as an internal standard. One µl aliquots of each sample were injected into a Shimadzu GC-17A gas chromatograph equipped with a Shimadzu QP5000 quadrupole electron ionization (EI) mass spectrometer (Shimadzu Scientific Instruments, Columbia, MD) as a detector. All analyses were made using splitless injections on a polar GC column (0.25mm diameter, 30m length, 0.25µm film thickness; EC WAX; W.R. Grace & Co., Columbia, MD) using ultra high purity (99.999%) helium as a mobile phase (split ratio 12:1, constant flow of 1ml/min). The gas chromatograph temperature and pressure parameters were optimized to resolve linalool and other floral volatiles to baseline with a total run time of 19 min per sample (240°C injection port temperature, 260°C detector temperature, 40°C initial temperature, two-min hold time increased at 15°C per min to 260°C, hold time 2.38 min). EI mass spectra (70 eV) were collected from m/z 40-350 (daltons) at a detector voltage of 70 eV, with scan speed of 1000 and a scan interval of 0.29 sec.

Volatile compounds were tentatively identified using computerized mass spectral libraries (Wiley Registry of Mass Spectral Data, National Institute of Standards and Technology, and [70] (> 120,000 mass spectra)). Peak areas were integrated using Shimadzu’s GCMSsolutions software and normalized for slight differences in final sample volume using the internal standard. Peak areas were quantified by comparison with five-point (log scale) dose-response standard curves generated using serial dilutions from 1 to 0.0001 mg/ml of authentic standards for (*E*)-β-ocimene, linalool, β-caryophyllene, geraniol, and (*E,E*)-farnesol. Emission rates were expressed as ng per flower, per hour (Table 1).

Additional chromatographic analyses were performed to determine which of the two known enantiomers of linalool is present in the floral headspace of *Oenothera harringtonii*. The third carbon of linalool (3,7-dimethylocta-1,6-dien-3-ol; C_10_H_18_O) is a chiral center, and both (S)-(+) and (R)-(-) enantiomers may be present in floral volatile blends [29, 71]. A chiral GC column (Cyclosil-B, Agilent Technologies, Inc.; 30m long, 0.25mm inner diameter, 0.25um film thickness) was used with the following temperature program: splitless injection at 240°C, column temperature increase from 40 to 240°C at 10°C per min. Under these conditions, the (R)-(-) and (S)-(+) enantiomers of linalool were distinguished by co-injection of two authentic standards, racemic linalool (Aldrich L260-2) and (R)-(-)-linalool (Aldrich 62139), along with a one-hour headspace sample taken from a single flower of *Clarkia breweri*, previously shown to exclusively emit (S)-(+) linalool [72]. Floral headspace samples from linalool+ populations of *O. harringtonii* (Florence) were chromatographed using the same protocol, comparing retention times for the leading edge of the linalool peak with those of the authentic standards.

### RNA Extraction

RNA was extracted from the same flowers used in fragrance analysis. While still attached to the plant, each flower was dissected into four parts: (1) stamens, (2) petals, (3) stigma and style, and (4) hypanthium and sepals. Additionally, RNA was extracted from flower buds 24-48 hours before anthesis and from young leaf tissues from the center of the rosette (excluding individuals D and F, two linalool-individuals for which bud and leaf samples could not be obtained due to insufficient growth post initial sampling). After tissues were excised from the plant, they were immediately placed into either 15ml conical tubes (petals, hypanthium, bud) or 2ml microcentrifuge tubes (stigma and style, stamens, leaf) and submerged in liquid nitrogen.

Frozen tissue was ground into a fine powder using either a mortar and pestle (petals, hypanthium, bud) or tissue homogenizer with a 3mm stainless steel ball bearing inside the microcentrifuge tube (stigma and style, stamens, leaf; Talboys High Throughput Homogenizer 930145, Troemner, Thorofare, NJ). RNA extraction was carried out with the Spectrum Plant Total RNA Mini Kit (Sigma-Aldrich, St. Louis, MO) or the RNeasy Mini Kit (Qiagen, Hilden, Germany), with an optional second elution to yield a total of 100μl of eluate. The amount of resulting RNA was quantified with the Qubit 2.0 fluorometer (Invitrogen, Carlsbad, CA) using the RNA Broad-Range Assay Kit (Invitrogen, Carlsbad, CA). All RNA samples were stored at − 80°C.

### Sequencing and Transcriptome Assembly

Sequencing of samples was conducted by the University of Chicago Genomics core using an Illumina HiSeq2500 (Illumina, San Diego, CA) with paired-end 100 base pair reads. All sequences were evaluated for quality using FastQC v0.10.1 [73]. Adapters and reads with low quality scores were trimmed from the readset with Trimmomatic v0.30 (ILLUMINACLIP:TruSeq3-PE.fa:2:30:10 LEADING:3 TRAILING:3 SLIDINGWINDOW:4:15 MINLEN:36) [74]. Reads that were likely rRNA or plastid RNA were removed by aligning all reads against a database of *Oenothera* plastid genes and *O. harringtonii* rRNA transcripts using bowtie2 v2.2.6 [75]. This database was created by combining an *Oenothera* plastid database (NCBI accession numbers EU262887, EU262889, EU262890, EU262891, KT881170, KT881173, KT881176) with a database of rRNA transcripts identified with RNAmmer v1.2 [76] from a pre-existing *O. harringtonii* seedling transcriptome and confirming with an NCBI BLAST search (blastn, nucleotide collection database). All reads were pooled together and normalized as a single readset using the in silico read normalization strategy from Trinity r2014-07-17 [77, 78] with a maximum coverage setting of 50. Reads were assembled using Trinity [77, 78] with default settings.

Assembly quality was evaluated using several different metrics. Percent of reads mapped to the assembly using bowtie2 v2.2.6 [75] was compared across all samples to ensure that reads mapped back evenly across all individuals and tissue types. Mean contig length, median contig length, and contig N50 were calculated using the “TrinityStats.pl” script provided with Trinity. Annotation statistics were calculated according to O’Neil and Emrich [79], including the number of hits and unique hits in a BLASTx and tBLASTn (v2.2.30) search to the *Arabidopsis thaliana* proteome, number of reciprocal best hits, average ortholog hit ratio, and average collapse factor.

#### Functional Annotation

Assembled transcripts were annotated for inferred function using the Trinotate r2014-07-08 pipeline [46], which generates an SQlite database containing several types of functional annotations. These include sequence homology comparisons from BLAST+ v2.2.30 [80] and Swiss-Prot [81], protein domain identification from HMMER v2.3.2 [82] and PFAM [83], protein signal peptide identification from signalP v4.1 [84], transmembrane domain prediction from TmHMM v2.0c [85] and comparisons to several functional annotation databases including eggNOG [86], GO [87], and KEGG [88]. Protein sequences for these comparisons were predicted using TransDecoder r2014-07-04 (https://github.com/TransDecoder/TransDecoder/wiki).

### Differential Expression Analysis

Differential expression analysis followed the protocol recommended by Trinity [77, 78] and scripts provided with Trinity; these scripts use R v3.3.0 [89] and several R packages (edgeR v3.12.0, Biobase v2.30.0, gplots v3.0.1, ape v3.4, qvalue v2.2.2, and goseq v1.22.0). The reads in each sample were mapped to the assembled contigs to determine gene expression levels using RSEM v1.2.15 [90]. In this analysis, there were six tissue types (petals; stigma and style; stamens; hypanthium and sepals; leaf; bud), two conditions (linalool+ and linalool-), and three biological replicates (excluding leaf and bud samples for linalool-, which only had one biological replicate).

Expression levels were calculated with the R package edgeR v3.12.0 [91, 92] to determine differentially expressed genes (DEGs). Pairwise comparisons between linalool+ and linalool-floral organs, among linalool-floral organs, and among linalool+ tissue types were conducted; a gene was considered differentially expressed if it had at least a four-fold change in expression and a p-value less than 0.001. This approach captures differences between floral organ expression and chemotype expression while minimizing the number of DEGs extracted that are not involved in floral fragrance biosynthesis. Clusters of DEGs with a similar expression pattern were created by cutting the DEG dendrogram at 65% of its height.

### GO Enrichment Analysis and Terpene Biosynthesis Gene Identification

Gene ontology (GO) enrichment analysis, performed with scripts provided in the Trinity software package, was used to identify gene categories that are overrepresented in each cluster relative to the assembly. Phylogenetic inference was used to identify and classify terpene synthases (TPS) in the reference transcriptome assembly. To classify putative terpene synthases into the seven described subfamilies a-f [23, 42, 93, 94], all of the sequences included in van Schie et al. [23] were downloaded and verified by reproducing a tree that circumscribed the seven TPS subfamilies. The verified alignment was then used to create a profile Hidden Markov Model using HMMER v3.1b (hmmer.org). The *O. harringtonii* reference assembly was searched for sequences explained by the pHMM using hmmscan (e-value 1e-10). All putative terpene synthesis protein sequences were aligned using MAFFT v7.310 [95] (--maxiterate 1000 -- localpair), and a phylogeny was inferred using RAxML v8.2.11 [96] with the PROTGAMMAWAG model of sequence evolution and 100 fast bootstrap replicates.

### SNP Analysis

To determine the distribution of SNPs in the putative (R)-(-)-linalool synthase coding sequence, a putative full length transcript was created by merging partial transcripts identified by the phylogenetic and DEG analyses. Three transcripts were merged that formed a clade (DN62538_c3_g1_i1, DN62538_c3_g2_i2, and DN74292_c2_g1_i11) and could be aligned with only a single mismatch. A web BLAST search against the GenBank database of non-redundant proteins resulted in alignment over approximately 52% of the length of (R)-(-)-linalool synthase from the lemon myrtle *Backhousia citriodora* (Myrtaceae), suggesting that the consensus sequence produced by merging the transcripts may not represent a full length transcript of (R)-(-)-linalool synthase. However, the merged *O. harringtonii* (R)-(-)-linalool synthase transcript and the lemon myrtle transcript (GenBank accession AB438045) were only 64% identical and it is possible that the full-length transcript in *Oenothera harringtonii* may be shorter. SNPs were then identified by mapping reads to the merged transcript from each RNA-Seq library. All reads identified in the DEG analysis as mapping to the three unmerged transcripts were extracted and variants were discovered using the GATK best practices workflow for germline SNPs (https://software.broadinstitute.org/gatk/best-practices/workflow?id=11145). Briefly, reads were re-mapped to the merged transcript and duplicate mappings were marked and removed before variants were called using HaplotypeCaller in GATK. Phasing of alleles within individuals was done with WhatsHap [97], after which two alternate phased haplotypes were extracted using bcftools (https://samtools.github.io/bcftools/). Each of the phased allele sequences, along with several linalool synthase accessions from the web BLAST search, were added to the previously generated alignment of terpene synthase sequences using the --add-alignment feature in MAFFT. A new gene family phylogeny was generated using RAxML as previously described.

## List of Abbreviations

DEG: Differentially expressed genes
FDR: False discovery rate
GC-MS: Gas chromatography-mass spectrometry
GO: Gene ontology
LIS: Linalool synthase
PC: Principal component
TPS: Terpene synthase
pHMM: profile Hidden Markov Model

## Declarations

### Ethics Approval and Consent to Participate

Not applicable.

### Consent for Publication

Not applicable.

### Availability of Data and Material

The Illumina RNA-Seq reads supporting the conclusions of this article are available from the NCBI Sequence Read Archive (BioProject PRJNA432694). The reference transcriptome assembly, mapping results, terpene synthase HMM and the alignment of terpene synthase sequences for phylogenetic analysis, and files used for the SNP analysis are currently available from the following link: https://doi.org/10.5061/dryad.np5hqbzrz.

### Authors’ Contributions

LLB, MGJ, GTB, RAL, RPO, TJ, JBF, RAR, KAS, and NJW designed the study, interpreted the data, and wrote the manuscript. LLB, RPO, GTB, and RAR performed the experiments. LLB, MGJ, GTB, TJ, RAR, and NJW analyzed the data. All authors read and approved the final manuscript.

## Acknowledgements

Thanks to Sandra Steiger (Justus-Liebig-Universität Gießen, Germany) for providing an authentic standard for methyl geranate. The authors thank Evan Hilpman, Sadie Todd, and Heather Rose Kates for assistance with floral fragrance collection in the field.

## Additional Files

Additional File 1: Number of raw and clean paired reads per sample, and SRA BioSample IDs. (XLS)

Additional File 2: All GO term enrichments for each cluster of differentially expressed genes. (XLS)

Additional File 3: GC-MS evidence for the absence of constitutive volatile terpenoid emissions from the leaves of *Oenothera harringtonii*. Panel A: Total ion chromatograms comparing floral headspace (upper trace, blue) with two samples taken from non-blooming vegetative rosettes (middle and lower traces, red and black) of *O. harringtonii* from a population (Florence, Colorado, USA) producing (R)-(-)-linalool in flowers. GC traces are presented at the same scale (T = internal standard of 23.6 ng toluene) and numbered compounds represent volatile terpenoids (1 = β-myrcene, 2 = (Z)-β-ocimene, 3 = (E)-β-ocimene, 4 = ocimene epoxide, L = (R)-(-)- linalool, 5 = β-caryophyllene, 6 = α-humulene). Note that all of these peaks (including 4) are absent in the vegetative samples. Panels B, C and D show the mass spectra (m/z 40-350) at 9.15 minutes, the retention time for linalool under the chromatographic conditions used in this study (see methods). The correct spectrum for linalool is present only in panel B, corresponding to the floral headspace; the two vegetative samples shown have no peak at this retention time. (PDF)

Additional File 4: Floral volatile data from parental field populations of *Oenothera harringtonii* used to set up transcriptomics experiments in greenhouse-grown plants. (XLS)

Additional File 5: Phylogeny of terpene synthases including phased alleles for the putative (R)-(-)-linalool synthase reconstructed in this study. (Newick)

## References

1. Shuttleworth A, Johnson SD. The missing stink: sulphur compounds can mediate a shift between fly and wasp pollination systems. Proc R Soc B Biol Sci. 2010;277:2811–9.

2. Raguso RA. Wake up and smell the roses: The ecology and evolution of floral scent. Annu Rev Ecol Evol Syst. 2008;39:549–69.

3. Junker RR, Blüthgen N. Floral scents repel facultative flower visitors, but attract obligate ones. Ann Bot. 2010;105:777–82.

4. Junker RR, Bluethgen N. Floral scents repel potentially nectar-thieving ants. Evol Ecol Res. 2008;10:295–308.

5. Kessler D, Gase K, Baldwin IT. Field experiments with transformed plants reveal the sense of floral scents. Science. 2008;321:1200–2.

6. Ashman T-L, Bradburn M, Cole DH, Blaney BH, Raguso RA. The scent of a male: The role of floral volatiles in pollination of a gender dimorphic plant. Ecology. 2005;86:2099–105.

7. Terry I, Walter GH, Moore C, Roemer R, Hull C. Odor-mediated push-pull pollination in cycads. Science. 2007;318:70.

8. Knudsen JT, Eriksson R, Gershenzon J, Ståhl B. Diversity and distribution of floral scent. Bot Rev. 2006;72:1–120.

9. Delle-Vedove R, Schatz B, Dufay M. Understanding intraspecific variation of floral scent in light of evolutionary ecology. Ann Bot. 2017;120:1–20.

10. Gross K, Sun M, Schiestl FP. Why do floral perfumes become different? Region-specific selection on floral scent in a terrestrial orchid. PLoS One. 2016;11:e0147975.

11. Sun M, Gross K, Schiestl FP. Floral adaptation to local pollinator guilds in a terrestrial orchid. Ann Bot. 2014;113:289–300.

12. Mant J, Peakall R, Schiestl FP. Does selection on floral odor promote differentiation among populations and species of the sexually deceptive orchid genus Ophrys? Evolution. 2005;59:1449–63.

13. Suinyuy TN, Donaldson JS, Johnson SD. Geographical variation in cone volatile composition among populations of the African cycad Encephalartos villosus. Biol J Linn Soc Lond. 2012;106:514–27.

14. Nagegowda DA. Plant volatile terpenoid metabolism: Biosynthetic genes, transcriptional regulation and subcellular compartmentation. FEBS Lett. 2010;584:2965–73.

15. Onda Y, Mochida K, Yoshida T, Sakurai T, Seymour RS, Umekawa Y, et al. Transcriptome analysis of thermogenic Arum concinnatum reveals the molecular components of floral scent production. Sci Rep. 2015;5:08753.

16. Magnard J-L, Roccia A, Caissard J-C, Vergne P, Sun P, Hecquet R, et al. Biosynthesis of monoterpene scent compounds in roses. Science. 2015;349:81–3.

17. Yue Y, Yu R, Fan Y. Transcriptome profiling provides new insights into the formation of floral scent in Hedychium coronarium. BMC Genomics. 2015;16:470.

18. Hsiao Y-Y, Tsai W-C, Kuoh C-S, Huang T-H, Wang H-C, Wu T-S, et al. Comparison of transcripts in Phalaenopsis bellina and Phalaenopsis equestris (Orchidaceae) flowers to deduce monoterpene biosynthesis pathway. BMC Plant Biol. 2006;6:14.

19. Amrad A, Moser M, Mandel T, de Vries M, Schuurink RC, Freitas L, et al. Gain and loss of floral scent production through changes in structural genes during pollinator-mediated speciation. Curr Biol. 2016;26:3303–12.

20. Wong DCJ, Amarasinghe R, Rodriguez-Delgado C, Eyles R, Pichersky E, Peakall R. Tissue-specific floral transcriptome analysis of the sexually deceptive orchid Chiloglottis trapeziformis provides insights into the biosynthesis and regulation of its unique UV-B dependent floral volatile, Chiloglottone 1. Front Plant Sci. 2017;8:1260.

21. Pichersky E, Raguso RA. Why do plants produce so many terpenoid compounds? New Phytol. 2016;3:692–702.

22. Chen F, Tholl D, Bohlmann J, Pichersky E. The family of terpene synthases in plants: A mid-size family of genes for specialized metabolism that is highly diversified throughout the kingdom. Plant J. 2011;66:212–29.

23. CCN van Schie, Haring MA, Schuurink RC. Tomato linalool synthase is induced in trichomes by jasmonic acid. Plant Mol Biol. 2007;64:251–63.

24. Linhart YB, Thompson JD. Thyme is of the essence: Biochemical polymorphism and multi-species deterrence. Evol Ecol Res. 1999;1:151–71.

25. Linhart YB, Gauthier P, Keefover-Ring K, Thompson JD. Variable phytotoxic effects of Thymus vulgaris (Lamiaceae) terpenes on associated species. Int J Plant Sci. 2015;176:20–30.

26. Gouyon PH, Vernet P, Guillerm JL, Valdeyron G. Polymorphisms and environment: The adaptive value of the oil polymorphisms in Thymus vulgaris L. Heredity. 1986;57:59–66.

27. Vernet P, Gouyon RH, Valdeyron G. Genetic control of the oil content in Thymus vulgaris L: A case of polymorphism in a biosynthetic chain. Genetica. 1986;69:227–31.

28. Wagner WL, Stockhouse RE, Klein WM. The systematics and evolution of the Oenothera caespitosa species complex (Onagraceae). Monographs in systematic botany from the Missouri Botanical Garden (USA). 1985;12: 1–103.

29. Raguso RA, Pichersky E. New Perspectives in Pollination Biology: Floral Fragrances. A day in the life of a linalool molecule: Chemical communication in a plant-pollinator system. Part 1: Linalool biosynthesis in flowering plants. Plant Species Biol. 1999;14:95–120.

30. Kawaano S, Odaki M, Yamaoka R, Oda-Tanabe M, Takeuchi M, Kawano N. Pollination biology of Oenothera (Onagraceae). The interplay between floral UV-absorbancy patterns and floral volatiles as signals to nocturnal insects. Plant Species Biol. 1995;10:31–8.

31. Raguso RA, Kelber A, Pfaff M, Levin RA, McDade LA. Floral biology of north american Oenothera sect. Lavauxia (Onagraceae): Advertisements, rewards, and extreme variation in floral depth. Ann Mo Bot Gard. 2007;94:236–57.

32. Boachon B, Junker RR, Miesch L, Bassard J-E, Höfer R, Caillieaudeaux R, et al. CYP76C1 (Cytochrome P450)-mediated linalool metabolism and the formation of volatile and soluble linalool oxides in Arabidopsis flowers: A strategy for defense against floral antagonists. Plant Cell. 2015;27:2972–90.

33. Reisenman CE, Riffell JA, Bernays EA, Hildebrand JG. Antagonistic effects of floral scent in an insect-plant interaction. Proc Biol Sci. 2010;277:2371–9.

34. Raguso RA. More lessons from linalool: Insights gained from a ubiquitous floral volatile. Curr Opin Plant Biol. 2016;32:31–6.

35. Okamoto T. Species-specific floral scents as olfactory cues in pollinator moths. In: Kato M, Kawakita A, editors. Obligate Pollination Mutualism. Tokyo: Springer Japan; 2017. p. 169–79.

36. Rhodes MK, Fant JB, Skogen KA. Local topography shapes fine-scale spatial genetic structure in the Arkansas Valley evening primrose, Oenothera harringtonii (Onagraceae). J Hered. 2014;105:806–15.

37. Bischoff M, Raguso RA, Jürgens A, Campbell DR. Context-dependent reproductive isolation mediated by floral scent and color. Evolution. 2015;69:1–13.

38. Kessler D, Kallenbach M, Diezel C, Rothe E, Murdock M, Baldwin IT. How scent and nectar influence floral antagonists and mutualists. Elife. 2015;4:e07641.

39. Bruzzese DJ, Wagner DL, Harrison T, Jogesh T, Overson RP, Wickett NJ, et al. Phylogeny, host use, and diversification in the moth family Momphidae (Lepidoptera: Gelechioidea). PLoS One. 2019;14:e0207833.

40. Artz DR, Villagra CA, Raguso RA. Spatiotemporal variation in the reproductive ecology of two parapatric subspecies of Oenothera cespitosa (Onagraceae). Am J Bot. 2010;97:1498–510.

41. He J, Fandino RA, Halitschke R, Luck K, Köllner TG, Murdock MH, et al. An unbiased approach elucidates variation in (S)-(+)-linalool, a context-specific mediator of a tri-trophic interaction in wild tobacco. Proc Natl Acad Sci USA. 2019;116:14651–60.

42. Dudareva N, Cseke L, Blanc VM, Pichersky E. Evolution of floral scent in Clarkia: novel patterns of S-linalool synthase gene expression in the C. breweri flower. Plant Cell. 1996;8:1137–48.

43. Cseke L, Dudareva N, Pichersky E. Structure and evolution of linalool synthase. Mol Biol Evol. 1998;15:1491–8.

44. Byng JW, Chase MW, Christenhusz MJM, Fay MF, Judd WS, Mabberley DJ, et al. An update of the Angiosperm Phylogeny Group classification for the orders and families of flowering plants: APG IV. Bot J Linn Soc. 2016;181:1–20.

45. Theiss KE, Holsinger KE, Evans MEK. Breeding system variation in 10 evening primroses (Oenothera sections Anogra and Kleinia; Onagraceae). Am J Bot. 2010;97:1031–9.

46. Bryant DM, Johnson K, DiTommaso T, Tickle T, Couger MB, Payzin-Dogru D, et al. A tissue-mapped axolotl de novo transcriptome enables identification of limb regeneration factors. Cell Rep. 2017;18:762–76.

47. Lamesch P, Berardini TZ, Li D, Swarbreck D, Wilks C, Sasidharan R, et al. The Arabidopsis Information Resource (TAIR): improved gene annotation and new tools. Nucleic Acids Res. 2012;40:D1202–10.

48. Ginglinger J-F, Boachon B, Höfer R, Paetz C, Köllner TG, Miesch L, et al. Gene coexpression analysis reveals complex metabolism of the monoterpene alcohol linalool in Arabidopsis flowers. Plant Cell. 2013;25:4640–57.

49. Jones CG, Moniodis J, Zulak KG, Scaffidi A, Plummer JA, Ghisalberti EL, et al. Sandalwood fragrance biosynthesis involves sesquiterpene synthases of both the terpene synthase (TPS)-a and TPS-b subfamilies, including santalene synthases. J Biol Chem. 2011;286:17445–54.

50. Huang M, Sanchez-Moreiras AM, Abel C, Sohrabi R, Lee S, Gershenzon J, et al. The major volatile organic compound emitted from Arabidopsis thaliana flowers, the sesquiterpene (E)-β-caryophyllene, is a defense against a bacterial pathogen. New Phytol. 2012;193:997–1008.

51. Kessler A, Baldwin IT. Defensive function of herbivore-induced plant volatile emissions in nature. Science. 2001;291:2141–4.

52. Xiao Y, Wang Q, Erb M, Turlings TCJ, Ge L, Hu L, et al. Specific herbivore-induced volatiles defend plants and determine insect community composition in the field. Ecol Lett. 2012;15:1130–9.

53. Huang X-Z, Chen J-Y, Xiao H-J, Xiao Y-T, Wu J, Wu J-X, et al. Dynamic transcriptome analysis and volatile profiling of Gossypium hirsutum in response to the cotton bollworm Helicoverpa armigera. Sci Rep. 2015;5:11867.

54. Schnee C, Köllner TG, Held M, Turlings TCJ, Gershenzon J, Degenhardt J. The products of a single maize sesquiterpene synthase form a volatile defense signal that attracts natural enemies of maize herbivores. Proc Natl Acad Sci USA. 2006;103:1129–34.

55. Clark EL, Carroll AL, Huber DPW. Differences in the constitutive terpene profile of lodgepole pine across a geographical range in British Columbia, and correlation with historical attack by mountain pine beetle. Can Entomol. 2010;142:557–73.

56. Pichersky E, Raguso RA, Lewinsohn E, Croteau R. Floral scent production in Clarkia (Onagraceae) (I. Localization and developmental modulation of monoterpene emission and linalool synthase activity). Plant Physiol. 1994;106:1533–40.

57. Raguso RA, Pichersky E. Floral volatiles from Clarkia breweri and C. concinna (Onagraceae): Recent evolution of floral scent and moth pollination. Plant Syst Evol. 1995;194:55–67.

58. Briscoe Runquist RD, Chu E, Iverson JL, Kopp JC, Moeller DA. Rapid evolution of reproductive isolation between incipient outcrossing and selfing Clarkia species. Evolution. 2014;68:2885–900.

59. Jia JW, Crock J, Lu S, Croteau R, Chen XY. (3R)-Linalool synthase from Artemisia annua L.: cDNA isolation, characterization, and wound induction. Arch Biochem Biophys. 1999;372:143–9.

60. Honaas LA, Wafula EK, Wickett NJ, Der JP, Zhang Y, Edger PP, et al. Selecting superior de novo transcriptome assemblies: Lessons learned by leveraging the best plant genome. PLoS One. 2016;11:e0146062.

61. Everaert C, Luypaert M, Maag JLV, Cheng QX, Dinger ME, Hellemans J, et al. Benchmarking of RNA-sequencing analysis workflows using whole-transcriptome RT-qPCR expression data. Sci Rep. 2017;7:1559.

62. Griffith M, Griffith OL, Mwenifumbo J, Goya R, Morrissy AS, Morin RD, et al. Alternative expression analysis by RNA sequencing. Nat Methods. 2010;7:843–7.

63. Shi Y, He M. Differential gene expression identified by RNA-Seq and qPCR in two sizes of pearl oyster (Pinctada fucata). Gene. 2014;538:313–22.

64. Huang M, Abel C, Sohrabi R, Petri J, Haupt I, Cosimano J, et al. Variation of herbivore-induced volatile terpenes among Arabidopsis ecotypes depends on allelic differences and subcellular targeting of two terpene synthases, TPS02 and TPS03. Plant Physiol. 2010;153:1293–310.

65. Xu Y-H, Wang J-W, Wang S, Wang J-Y, Chen X-Y. Characterization of GaWRKY1, a cotton transcription factor that regulates the sesquiterpene synthase gene (+)-delta-cadinene synthase-A. Plant Physiol. 2004;135:507–15.

66. Nieuwenhuizen NJ, Chen X, Wang MY, Matich AJ, Perez RL, Allan AC, et al. Natural variation in monoterpene synthesis in kiwifruit: transcriptional regulation of terpene synthases by NAC and ETHYLENE-INSENSITIVE3-like transcription factors. Plant Physiol. 2015;167:1243–58.

67. Singh N, Sharma A. In-silico identification of miRNAs and their regulating target functions in Ocimum basilicum. Gene. 2014;552:277–82.

68. Sobhani Najafabadi A, Naghavi MR. Mining Ferula gummosa transcriptome to identify miRNAs involved in the regulation and biosynthesis of terpenes. Gene. 2018;645:41–7.

69. Skogen KA, Jogesh T, Hilpman ET, Todd SL, Rhodes MK, Still SM, et al. Land-use change has no detectable effect on reproduction of a disturbance-adapted, hawkmoth-pollinated plant species. Am J Bot. 2016;103:1950–63.

70. Adams RP. Identification of essential oil components by gas chromatography/quadrupole mass spectroscopy. 3rd ed. Illinois: Allured Publishing Corporation; 2001:456pp.

71. Hanneguelle S, Thibault JN, Naulet N, Martin GJ. Authentication of essential oils containing linalool and linalyl acetate by isotopic methods. J Agric Food Chem. 1992;40:81–7.

72. Pichersky E, Lewinsohn E, Croteau R. Purification and characterization of S-linalool synthase, an enzyme involved in the production of floral scent in Clarkia breweri. Arch Biochem Biophys. 1995;316:803–7.

73. Andrews S. FastQC: A quality control tool for high throughput sequence data [Online]. 2010. https://www.bioinformatics.babraham.ac.uk/projects/fastqc/.

74. Bolger AM, Lohse M, Usadel B. Trimmomatic: A flexible trimmer for Illumina sequence data. Bioinformatics. 2014;30(i15):2114–20.

75. Langmead B, Salzberg SL. Fast gapped-read alignment with Bowtie 2. Nat Methods. 2012;9:357–9.

76. Lagesen K, Hallin P, Rødland EA, Staerfeldt H-H, Rognes T, Ussery DW. RNAmmer: Consistent and rapid annotation of ribosomal RNA genes. Nucleic Acids Res. 2007;35:3100–8.

77. Grabherr MG, Haas BJ, Yassour M, Levin JZ, Thompson DA, Amit I, et al. Trinity: Reconstructing a full-length transcriptome without a genome from RNA-Seq data. Nat Biotechnol. 2011;29:644–52.

78. Haas BJ, Papanicolaou A, Yassour M, Grabherr M, Blood PD, Bowden J, et al. De novo transcript sequence reconstruction from RNA-seq using the Trinity platform for reference generation and analysis. Nat Protoc. 2013;8(8):1494–512.

79. O’Neil ST, Emrich SJ. Assessing De Novo transcriptome assembly metrics for consistency and utility. BMC Genomics. 2013;14:465.

80. Camacho C, Coulouris G, Avagyan V, Ma N, Papadopoulos J, Bealer K, et al. BLAST+: Architecture and applications. BMC Bioinformatics. 2009;10:421.

81. Boeckmann B, Blatter M-C, Famiglietti L, Hinz U, Lane L, Roechert B, et al. Protein variety and functional diversity: Swiss-Prot annotation in its biological context. C R Biol. 2005;328:882–99.

82. Eddy SR, Wheeler TJ. HMMER: Biosequence analysis using profile Hidden Markov Models. http://hmmer.org/.

83. Finn RD, Coggill P, Eberhardt RY, Eddy SR, Mistry J, Mitchell AL, et al. The Pfam protein families database: Towards a more sustainable future. Nucleic Acids Res. 2016;44:D279–85.

84. Petersen TN, Brunak S, von Heijne G, Nielsen H. SignalP 4.0: Discriminating signal peptides from transmembrane regions. Nat Methods. 2011;8(10):785–6.

85. Krogh A, Larsson B, von Heijne G, Sonnhammer EL. Predicting transmembrane protein topology with a hidden Markov model: Application to complete genomes. J Mol Biol. 2001;305:567–80.

86. Huerta-Cepas J, Szklarczyk D, Forslund K, Cook H, Heller D, Walter MC, et al. eggNOG 4.5: A hierarchical orthology framework with improved functional annotations for eukaryotic, prokaryotic and viral sequences. Nucleic Acids Res. 2016;44:D286–93.

87. Gene Ontology Consortium. Gene Ontology Consortium: going forward. Nucleic Acids Res. 2015;43:D1049–56.

88. Kanehisa M, Sato Y, Kawashima M, Furumichi M, Tanabe M. KEGG as a reference resource for gene and protein annotation. Nucleic Acids Res. 2016;44:D457–62.

89. R Core Team. R: A Language and Environment for Statistical Computing. https://www.R-project.org.

90. Li B, Dewey CN. RSEM: Accurate transcript quantification from RNA-Seq data with or without a reference genome. BMC Bioinformatics. 2011;12:323.

91. Robinson MD, McCarthy DJ, Smyth GK. edgeR: A Bioconductor package for differential expression analysis of digital gene expression data. Bioinformatics. 2010;26:139–40.

92. McCarthy DJ, Chen Y, Smyth GK. Differential expression analysis of multifactor RNA-Seq experiments with respect to biological variation. Nucleic Acids Res. 2012;40:4288–97.

93. Bohlmann J, Steele CL, Croteau R. Monoterpene synthases from grand fir (Abies grandis). cDNA isolation, characterization, and functional expression of myrcene synthase, (-)-(4S)-limonene synthase, and (-)-(1S,5S)-pinene synthase. J Biol Chem. 1997;272:21784–92.

94. Dudareva N, Martin D, Kish CM, Kolosova N, Gorenstein N, Fäldt J, et al. (E)-beta-ocimene and myrcene synthase genes of floral scent biosynthesis in snapdragon: function and expression of three terpene synthase genes of a new terpene synthase subfamily. Plant Cell. 2003;15:1227–41.

95. Katoh K, Standley DM. MAFFT multiple sequence alignment software version 7: Improvements in performance and usability. Mol Biol Evol. 2013;30:772–80.

96. Stamatakis A. RAxML version 8: A tool for phylogenetic analysis and post-analysis of large phylogenies. Bioinformatics. 2014;30:1312–3.

97. Patterson M, Marschall T, Pisanti N, van Iersel L, Stougie L, Klau GW, et al. WhatsHap: Weighted haplotype assembly for future-generation sequencing reads. J Comput Biol. 2015;22:498–509.

